# A new class of protein sensor links spirochete pleomorphism, persistence, and chemotaxis

**DOI:** 10.1101/2022.01.11.475842

**Authors:** A.R. Muok, K. Kurniyati, D.R. Ortega, F.A. Olsthoorn, A. Sidi Mabrouk, C. Li, A. Briegel

**Affiliations:** Institute of Biology, Leiden University, Leiden, The Netherlands; Centre for Microbial Cell Biology, Leiden University, Leiden, Netherlands; Department of Oral and Craniofacial Molecular Biology, Philips Research Institute for Oral Health, Virginia Commonwealth University, Richmond, VA, 23298, USA

## Abstract

Pathogenic spirochetes can alter their morphologies and behaviors to infect and survive within their hosts. Previous reports demonstrate that the formation of so-called ‘round bodies’ and biofilms, and chemotaxis are involved in spirochete pathogenesis. Here, in the spirochete *Treponema denticola*, we report a direct link between these cellular states that involves a new class of protein sensor (CheWS) with hitherto unclear function. Using cryo-EM methods, protein modeling, bioinformatics, genetics methods, and behavioral assays we demonstrate that spirochetes regulate these behaviors in response to the small molecule s-adenosylmethionine (SAM) via a SAM sensor that is anchored to chemotaxis arrays. CheWS influences chemotaxis, biofilm and round body formation under nonstressed conditions by a novel sporulation-like mechanism. Taken together, we establish an improved model for round body formation, we discovered a direct link between this SAM sensor and changes in cellular states, as well as characterized a new sensor class involved in chemotaxis.

## Introduction

Sensory systems allow microbes to adapt to and thrive in their environment. By sensing stimuli through ligand-specific proteins, bacteria can find nutrients and avoid toxic moieties, evade predators, adapt to their hosts or symbionts, and restructure their micro-environment. During these processes, bacteria often change their physiologies accordingly. For free-living bacteria, they allow the cells to find sources of food and withstand ecological changes^1^. For bacterial pathogens, these systems can be critical for infecting hosts tissues and sustaining infections^2,3^. Although numerous strides have been made in understanding how the underlying molecular networks that facilitate these sensory systems operate, many key players or mechanisms remain unclear.

For many bacteria, the appropriate response to some stimuli is to enter a non-motile cellular state. For example, when the environment is unfavorable, bacteria can form non-motile pleomorphic states and biofilms that protect the cells from external stresses^4–6^. Pleomorphism describes the ability of microbes to alter their morphology in response to external conditions. When cells form biofilms, they cluster together and embed themselves in an extracellular matrix that offers physical protection from the environment. In pathogenic spirochetes such as *Treponema denticola* (*Td*), a causative agent of periodontal diseases^2^ and linked to Alzheimer’s disease^7,8^ and specific cancers^9^, it is established that specific pleomorphic states and biofilm formation are involved in host colonization and infection persistence (Fig. 1A)^5,10^. These pleomorphic states are called round bodies, a form of non-motile cell with minimized metabolic activity that are specific to spirochetes^10^. These structures form as a response to stress to physically protect cells from the environment, and a single round body may contain several spirochete cells^4,6,11^. These cells remain in a dormant state until the environment becomes favorable again and the cells can resume normal growth and behaviors^12^. Round body formation can be induced by a wide range of stresses including starvation^4^, hypotonic treatment^13,14^, osmotic pressure^6^, growth to stationary phase^15^, and antibiotic exposure^15^. In fact, round bodies can withstand otherwise lethal doses of some antibiotics^15,16^, which may account for recurrence of spirochetal infections after antibiotic treatments^17^. Round bodies are also the dominant cellular form in mature spirochete biofilms^17^, spinal fluid^12^, and brain tissues from Alzheimer’s patients^18^. In addition to offering stress protection, spirochete biofilms are also formed in response to interactions with microbial symbionts and host molecules^19,20^.

**Fig. 1:**
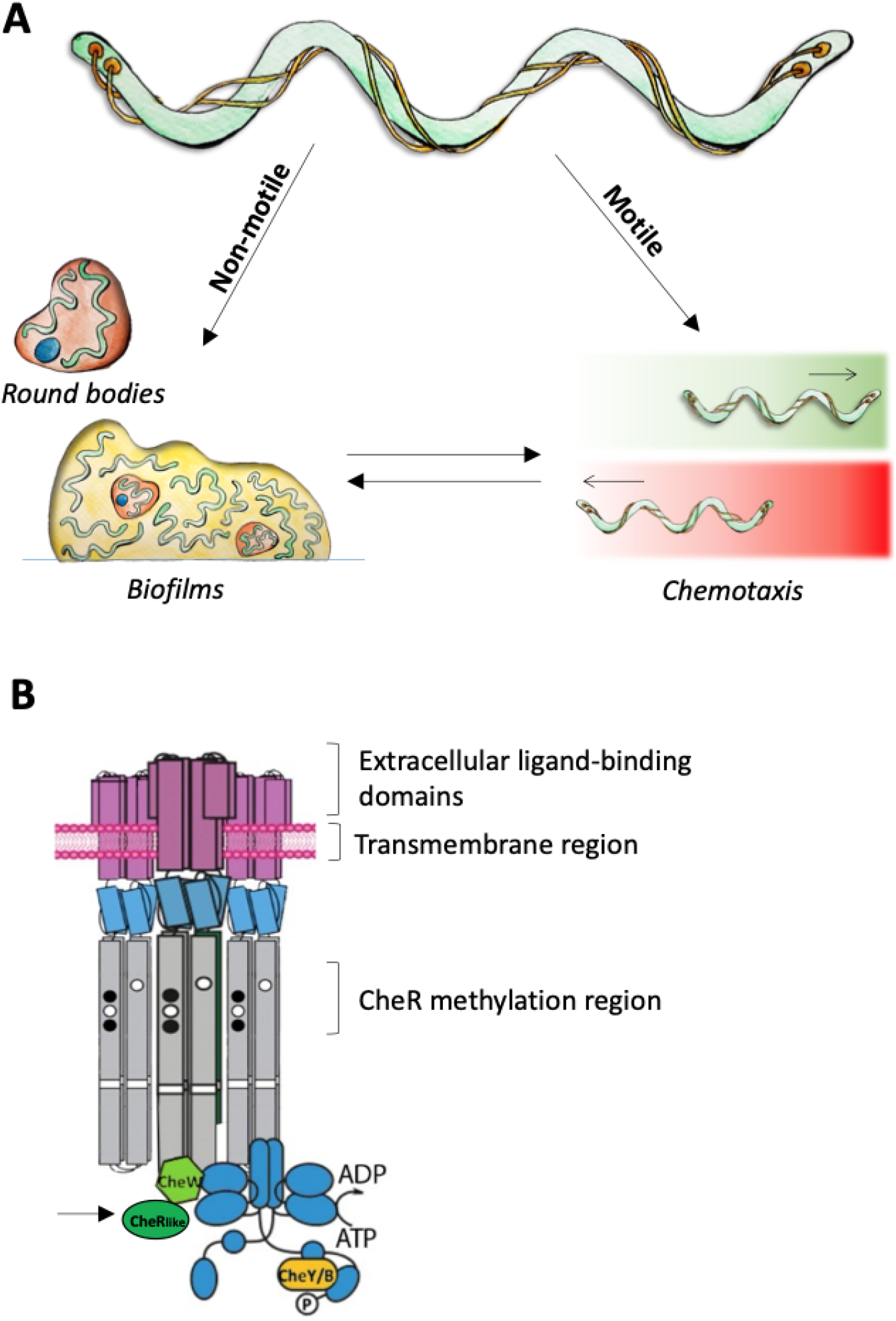
The life-cycle and chemotaxis machinery of spirochetes. **(A)** Spirochetes can form non-motile states that protect them from stress and contribute to host colonization. Motile spirochetes can undergo chemotaxis that enables them to move toward favorable environments (green gradient) and away from deleterious ones (red gradient). (B) The chemotaxis system consists of three main proteins: chemoreceptors that span the inner membrane, the kinase CheA (blue), and the adaptor protein CheW (green). *Td* CheW possesses an additional C-terminal domain, called the CheR_like_ domain (black arrow), that is a distant homolog of the CheR chemotaxis receptor methyltransferase.

It is unclear how signals switch spirochetes from motile to non-motile forms. In other organisms, like *Comamonas testosteroni*, it is established that the chemotaxis system is directly involved in transitions from motile states to sessile biofilm states^21^. Bacterial chemotaxis is the system that allows cells to sense their external environment through transmembrane chemoreceptors to ultimately control the motility apparatus of the cells, such as the flagella^22,23^. The chemoreceptors recognize specific chemicals with their extracellular domains and transduce signals into the cell. Two proteins, called CheA and CheW, bind to the intracellular tips of the receptors and initiate a phosphor-relay system when activated by the receptors (Fig. 1B)^24^. CheA is a five-domain histidine kinase that directly activates its response regulators via phosphoryl-transfer^25^. The protein CheW links CheA to receptors and collectively all three proteins form a membrane-associated hexagonal apparatus called the chemotaxis array. The initiated phosphor-relay subsequently controls the rotation direction of flagella, allowing the cell to move toward favorable environments and away from deleterious ones. It is established that chemotaxis is essential for pathogenesis of some spirochetes. For example, *in vivo* experiments show that the chemotaxis system in *Borrelia burgdorferi* (*Bb*), the causative agent for Lyme disease, is essential for infecting mice^3,26,27^.

In *Td*, several chemotaxis proteins are essential for penetrating oral epithelial cells^2^, forming monospecies biofilms^19^, and forming synergistic co-biofilms^19,20^. Furthermore, it has been recently shown that *Td* cells can actively move from oral cavities to brain tissues where they induce structures associated with Alzheimer’s disease in mice^28^, although the role of chemotaxis in this process is unknown. Recent research with *Td* has revealed that the structure of the chemotaxis system in spirochetes differs from canonical systems in several regards^29^. Their chemotaxis arrays have unique linear symmetry and contain a novel CheW variant that possesses an additional C-terminal domain. Previous findings show that this additional domain, called CheR_like_, is widespread among the spirochaetes (Fig. 1B)^29,30^. It is a distant homolog of the chemotaxis protein CheR, which methylates receptors using the methyl-donor S-adenosyl methionine (SAM) as part of an adaptation response^31,32^. In addition to a role in the chemotaxis system, SAM is a critical cofactor in methyl-transfer reactions that are essential for cell viability, both in eukaryotes and prokaryotes^33^. In prokaryotes specifically, SAM is essential for synthesis of some amino acids and may be important for microbiome interactions^34,35^. CheR_like_ can bind SAM but does not possess the conserved residues for enzyme catalysis or receptor binding^29,31,36^. Thus, the exact cellular role of this domain remained unclear.

Here, we applied cryo-electron tomography (cryo-ET) of *Td* cells to generate an improved model for spirochete round body formation. Namely, we show that log-phase *Td* cells form round bodies in a process similar to sporulation. This model suggests that spirochetes are capable of forming round bodies under non-stressed conditions, thus indicating higher virulence capabilities than previously thought. Furthermore, we show that the CheR_like_ domain influences round body and biofilm formation, and functions as a chemotaxis sensor for SAM. This finding, exemplified in *Td*, links transition from motile chemotactic cells to non-motile round bodies and biofilms. This is the first reported instance of a characterized CheW-fused chemotaxis sensor, which behests a new designation for this protein, that we have called CheWS (CheW ŞAM-sensor).

## Results

### Round body formation in *T. denticola*

Spirochetes are typically present as long, thin spiral-shaped cells that are motile and chemotactic, but can form round bodies and biofilms. Round bodies have been investigated in the spirochetes *Td*^37^, *Bb*^4,5,38^ and *Treponema pallidum* (*Tp*)^39^ through microscopy methods and has led to an established model for their formation (Fig. S1)^10^. This process generally occurs in two steps. First the spiral-shaped cell significantly enlarges its outer membrane at the cell tip so that the cell can then move into that newly-formed space. Once the cell is inside this protective membrane, it can divide to form typical spiral-shaped cells. Additionally, the round bodies may also contain dense, spherical cells called ‘core structures’ that are typically dominant in aged round bodies^13^. Here, the encased bacteria can ‘hide’ from the unfavorable environment and avoid cell death. When these round bodies are placed back into favorable, non-stressful environments, they form motile cells within a few days^5^. However, the core structures may take several weeks before forming the spiral-shaped, motile cells^5,12,13^. The ability of spirochetes to form round bodies has been described as a form of pleomorphism, as the cells are changing their morphology and metabolism in response to their environment.

These previous investigations of the round body formation were limited to traditional EM and light microscopy methods. As such, structural insights into this process has been lacking. Therefore, we conducted cryo-EM and cryo-ET experiments with *Td* WT cells undergoing round-body formation. We observed notable differences between cells in log phase and stationary phase growth. During the stationary phase, our images match the established model for round body formation (Fig. 2A). Large round bodies (~1-2 μm)^37^ are present and the typical spiral-shaped spirochete cells are clearly visible inside the round bodies (Fig. 2A,B). Several of these round bodies contain more than one cell, including core structures observed in other spirochetes (Fig. 2B). However, during log phase, we see much smaller (200-400 nm), densely-packed round bodies that form at the cell tips (Fig. 2C). These round bodies ultimately pinch off from the mother cell. In the early stages of this process, both the outer and inner membranes of the mother cell and round body are connected. At later stages, only the outer membrane is connected, which then separates to form a free round body and an intact mother cell (Fig. 2C). Unlike the established model, we neither see enlarged vacant outer membranes nor spiral cells inside the round body at any stage. Instead, the round bodies resemble the previously described electron dense core structures^12,13^. Therefore, in the log-phase growth, round body formation has a stronger resemblance to a sporulation process rather than pleomorphism. Collectively, these results were used to generate a more comprehensive model for round body formation that now includes log phase characterizations (Fig. 2D).

**Fig. 2:**
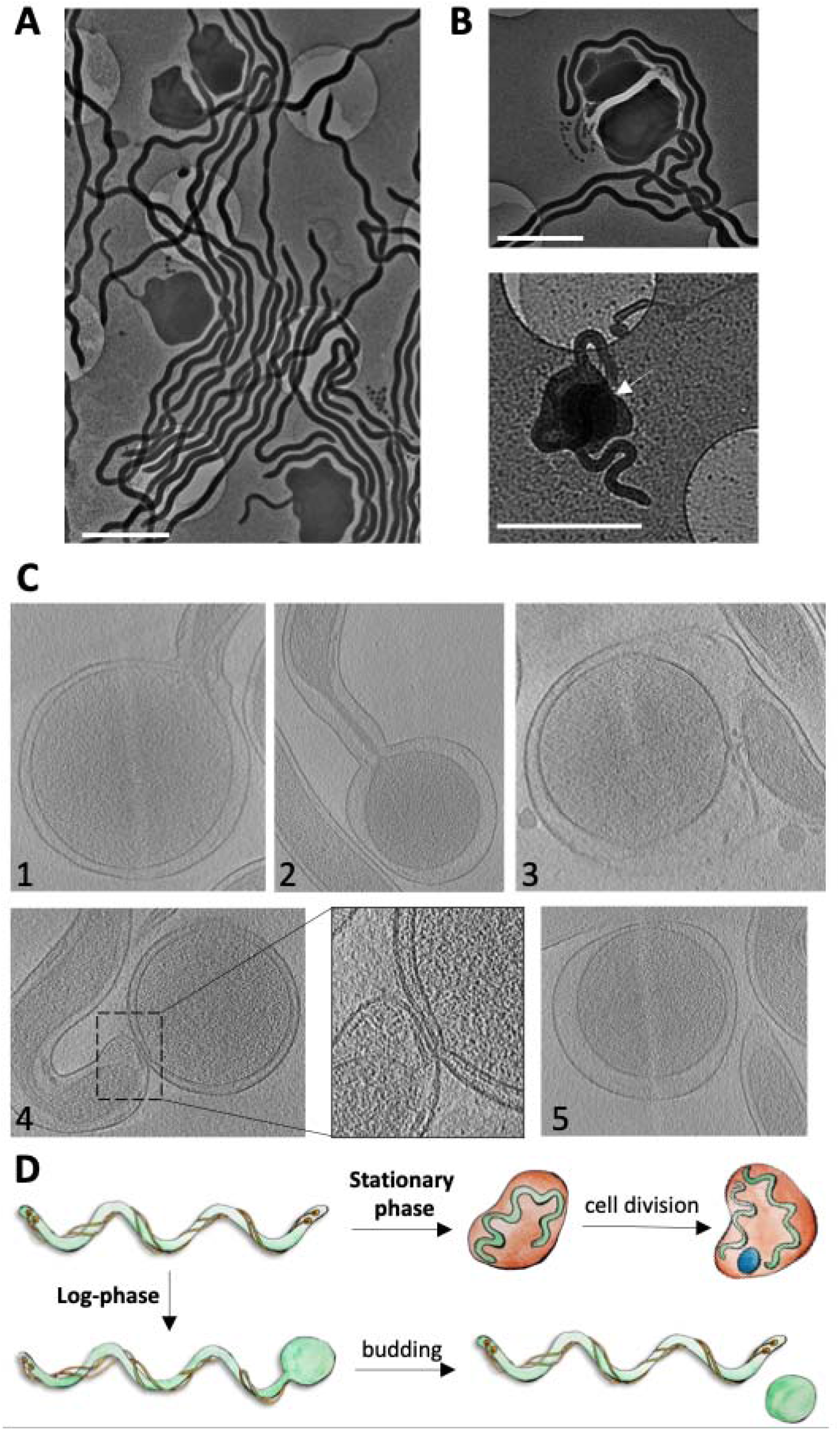
Cryo-electron microscopy reveals the morphology of *Td* round bodies. **(A)** In stationary phase, *Td* forms round bodies that resemble morphologies seen in previous reports. Scale bars are 2 μm. (B) The stationary phase round bodies may contain more than one spiral-shaped spirochete (top), or ‘core structure’ (white arrow) (bottom). Scale bars are 2 μm. (C) During log-phase, round bodies do not contain spiral-shaped cells, are smaller than stationary phase round bodies, and bud from the tips of spiral-shaped cells. The formation of these round bodies are shown in steps 1-5. 1: The outer and inner membrane of the cell tips begin to expand. 2: The inner membrane begins to pinch off from the forming round body. 3: The inner membrane of the round body and spiral cell are no longer connected but they are still attached via the outer membrane. 4: The outer membrane of the round body starts to disconnect from the spiral cell. 5: The round body is now separated from the spiral cell. (D) Log phase round body characterizations generate an improved model for round body formation, which previously only included stationary phase characterizations.

The log-phase round bodies possess two membranes and a continuous peptidoglycan layer during all stages of growth and after separation from the mother cell (Fig. S2). Additionally, some round bodies possess flagella and flagellar motors between the outer and inner membranes (Fig. S3). The round bodies do not contain visible chemoreceptor arrays. However, in some instances, arrays are visible within the mother cell in regions adjacent to the forming round body (Fig. S3).

### The CheR_like_ domain of CheWS influences round body formation

Previous cryo-ET experiments of motile (spiral-shaped) WT and ΔCheR_like_ cells have revealed a significant difference in the structural stabilization of chemotaxis arrays between the two strains^29^. Here, through cryo-EM we also observed a significant difference in the fraction of log phase cells that are in the process of forming round bodies. For the WT and ΔCheR_like_ strains, ~11.6 % and ~23.2 % of the cells are observed forming round bodies, respectively (Fig. 3A). The addition of 0.5 mM SAM almost completely eliminates observed round body formation in both strains, with no significant difference found between them (Fig. 3A). These results are summarized and compared in Supplementary Figure 4. These data show no discernible phenotypic difference between the two strains (Fig. 2C, 3B).

**Fig. 3:**
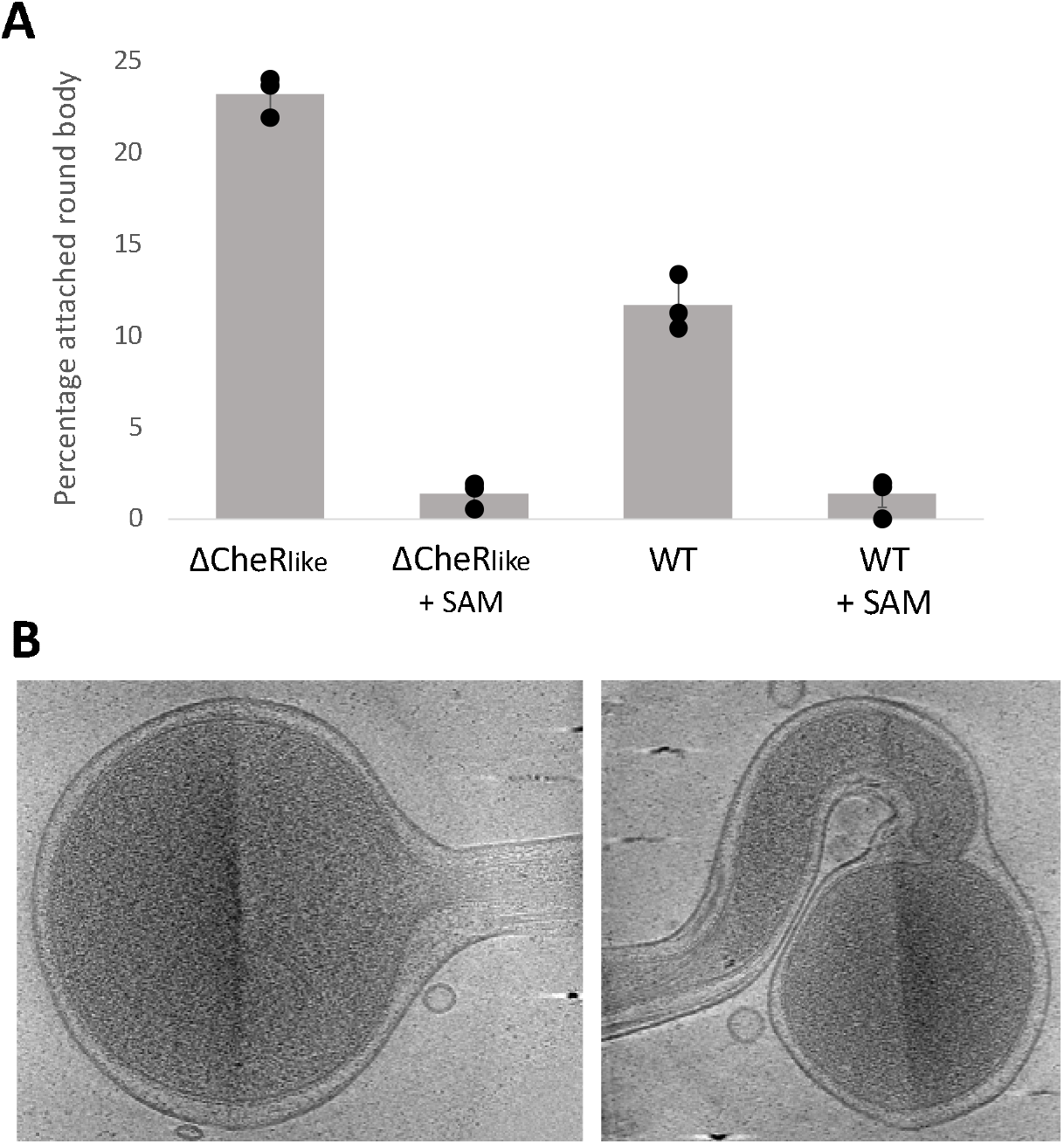
*Td* round body formation is influenced by the CheR_like_ domain. **(A)** The percentage of cells undergoing log phase round body formation were quantified in *Td* WT and a *Td* ΔCheR_like_ strain. The WT and ΔCheR_like_ strains have ~11.6 +/-1.51 % and ~23.2 +/-1.1 % of cells undergoing round body formation, respectively. There is a significant difference between the two strains using a two-tailed null hypothesis significance test (p < 0.05). When 0.5 mM SAM is added to the cells, the WT and ΔCheR_like_ strains have ~1.2 +/-1.1 % and ~1.4 +/- 0.7 % of cells undergoing round body formation, respectively, and there is no significant difference between the two strains (p > 0.05). However, for both strains, the presence of SAM significantly reduces the percentage of cells undergoing round body formation (p < 0.05). Results are expressed as the mean of cell percentages undergoing round body formation for three samples +/- one standard deviation. **(B)** Log phase round bodies in the ΔCheR_like_ strain are morphologically similar to the WT strain.

### The CheR_like_ domain of CheWS influences biofilm formation

As the ΔCheR_like_ strain has a significant increase in cells undergoing round body formation, we conducted assays to determine the subsequent impact on biofilm formation. Quantitative absorbance-based biofilm assays with the WT and ΔCheR_like_ strains reveal that the ΔCheR_like_ strain produces significantly less biofilm than the WT strain (Fig. 4A). The addition of 0.5 mM SAM significantly increases biofilm formation in both strains, but the ΔCheR_like_ Strain still forms significantly less biofilms than WT (Fig. 4A). These results are summarized and compared in Supplementary Figure 4.

**Fig. 4:**
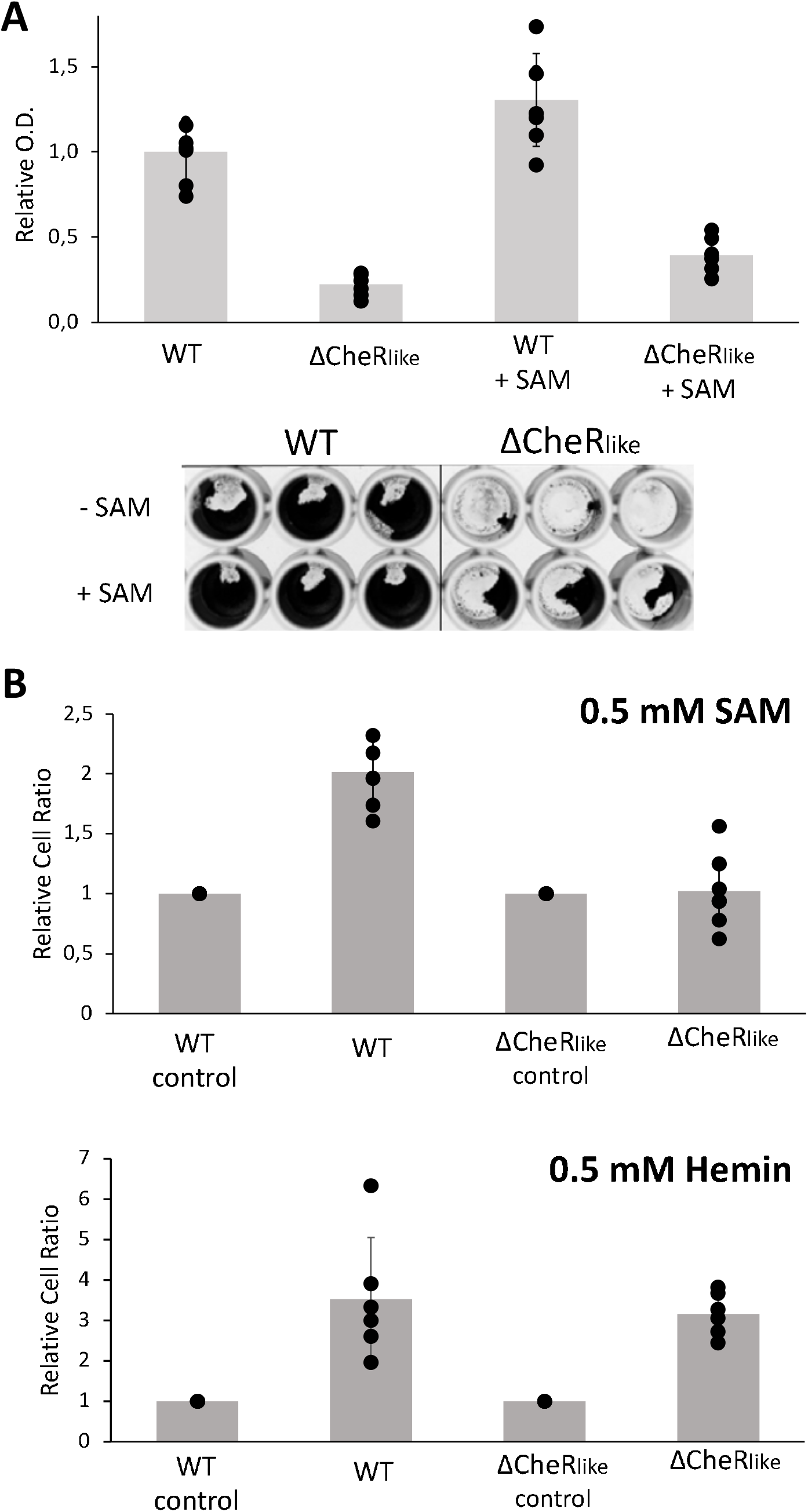
The CheR_like_ domain influences biofilm formation and chemotaxis toward SAM. **(A)** Quantitative biofilm assays show that the CheR_like_ domain significantly alters biofilm formation. The relative ratio of biofilms formed for the WT and ΔCheR_like_ strains are 1.0 +/- 0.17 and 0.22 +/- 0.06, respectively. There is a significant difference between the two strains using a two-tailed null hypothesis significance test (p < 0.05). When 0.5 mM SAM is added to the samples, the amount of relative biofilm for the WT and ΔCheR_like_ strains are 1.30 +/- 0.27 and 0.39 +/- 0.10. There is a significant difference in biofilm formation between the two strains in the presence of SAM (p < 0.05). Furthermore, for both strains, the addition of SAM significantly increases the relative amount of biofilm (p < 0.05). Results are normalized so that the mean for the WT strain without SAM added is equal to 1. Results are expressed as the mean OD_595_ for seven samples +/- one standard deviation. The bottom image is a representative experiment for relative biofilm quantification. (B) Chemotaxis assays demonstrate that the CheR_like_ domain is responsible for chemotaxis toward SAM (top) but not the previously established chemoattractant hemin (bottom). For the SAM capillary assays, the WT and ΔCheR_like_ strains have a relative ratio of 2.02 +/- 0.28 and 1.03 +/- 0.27 cells in the capillary tube compared to non-gradient controls, respectively. The WT strain is significantly different than nongradient controls using a two-tailed null hypothesis significance test (p < 0.05), but the ΔCheR_like_ strain is not significantly different than non-gradient controls (p > 0.05). For the hemin capillary assays, the WT and ΔCheR_like_ Strains have a relative ratio of 3.52 +/- 1.52 and 3.17 +/- 0.53 cells in the capillary tube compared to non-gradient controls, respectively. Both strains are significantly different than non-gradient controls (p < 0.05). Results are normalized so that the non-gradient controls are equal to 1 for all samples. Results are expressed as the mean cell count for all samples +/- one standard deviation. The SAM experiments were conducted with ten samples, and the hemin experiments were conducted with six samples.

### The CheR_like_ domain of CheWS modulates chemoattraction towards SAM

Previous research shows that the CheR_like_ domain does not possess the conserved residues for catalysis or receptor binding, but that it does bind SAM^29^. Therefore, we conducted capillary-based chemotaxis assays to determine if the CheR_like_ domain functions as a SAM sensor. While it was previously unknown if *Td* chemotactically responds to SAM, capillary assays with the WT strain demonstrate that it is a chemoattractant. However, the ΔCheR_like_ strain does not respond to SAM and the response is not significantly different from non-gradient negative controls (Fig. 4B). As a positive control, chemoattraction toward the established attractant hemin was also tested^40^. Both the wild-type (WT) and ΔCheR_like_ strains are attracted to hemin, with no significance difference found between the two strains (Fig. 4B). These results are summarized and compared in Supplementary Figure 4.

### Structural analysis of the CheR_like_ domain from *T. denticola* CheWS

Crystallography experiments of the CheR_like_ domain (residues 203-444) were unsuccessful, with no visible crystals forming in any tested conditions. When the CheR_like_ domain is expressed and purified without the N-terminal sub-domain (residues 258-444), the protein precipitates at ambient temperatures, indicating that smaller N-terminal sub-domain is critical for thermodynamic stability of the C-terminal sub-domain. The addition of the substrate SAM did not aid in the stabilization of the isolated C-terminal sub-domain.

A model of the *Td* CheR_like_ domain from the CheWS protein (residues 203-444) was determined using the automated AlphaFold 2.0 software^41^. The software produced five models that slightly vary in the Local Difference Distance Test (IDDT) and Predicted Align Error (PAE) scores (Fig. S5). We further analyzed the highest scored model (model 1). Like classical CheR proteins, the CheR_like_ model is a two-domain, mixed α/β protein (Fig. 5A)^31,42^. The N-terminal domain consists of three helices and is much smaller than the mixed C-terminal domain. The C-terminal domain possess an obvious cavity located at the expected the SAM-binding region. When we included the linker connecting the CheW and CheR_like_ domains (residues 175-202) in the AlphaFold 2.0 run, the structure of the linker could not be ascertained, likely indicating a highly flexible and disordered structure. We also generated an AlphaFold 2.0 model of the classical CheR protein in *Td*, which also does not have an experimentally derived structure (Fig. S6A).

**Fig 5:**
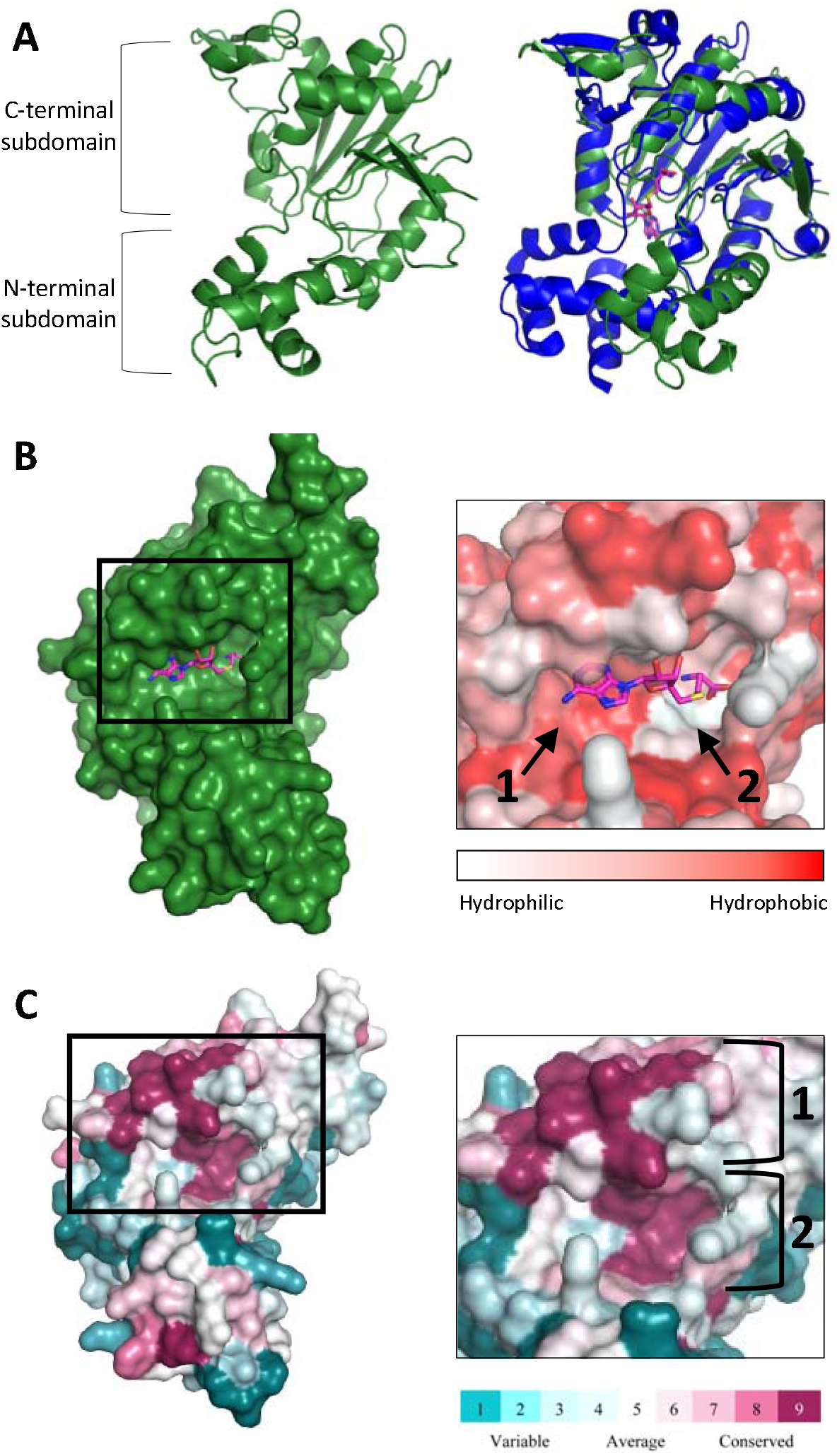
Structural modeling and conservation analysis of the *Td* CheR_like_ domain. **(A)** The CheR_like_ domain model (green) possesses the N-terminal and C-terminal sub-domains of classical CheR proteins (left). However, when compared to known structures of classical CheR proteins (blue, PDBID: 5FTW), the relative position of the N-terminal sub-domain is shifted (right). **(B)** These observed structural differences in the CheR_like_ domain model exposes the hypothetical SAM pocket to solution (left). The hypothetical SAM pocket is relatively hydrophobic near the adenine ring of SAH (1) but relatively polar near the homocysteine portion (2)(right). **(C)** Conservation analysis of the CheR_like_ domain model shows high residue conservation at a lid above the SAM pocket (1) and inside the SAM pocket (2).

While the CheR_like_ domain possesses the typical topology of classical CheR proteins, there are differences in tertiary interactions that result in a relative displacement of the N-terminal subdomain (Fig. 5A). Specifically, in the CheR_like_ domain, several loops of the N-terminal sub-domain interact with loops of the central beta-sheet in the C-terminal domain through hydrophobic contacts. The relative position of the N-terminal sub-domain of CheR_like_ exposes the expected SAM-binding pocket to the solution (Fig. 5A,B). In known classical CheR structures, and our *Td* classical CheR model, the substrate is buried in a cavity that is produced by interactions between the N- and C-terminal sub-domains (Fig. 5A)^31^. When the closest-related classical CheR homolog of known structure (*Bacillus subtilis* CheR with the SAM-analog, S-adenosyl homocysteine (SAH), PDBID: 5FTW)^42^ is aligned to the CheR_like_ structure, the SAH molecule in 5FTW fits perfectly within the expected substrate pocket of CheR_like_(Fig. 5B). Residue mapping using the Eiseberg hydrophobicity scale (PyMol) reveals that the CheR_like_ substrate pocket has high hydrophobicity near the adenine ring but is highly polar near the homocysteine portion of the molecule, which is also seen in known classical CheR structures (Fig. 5B)^31,36,42^.

To investigate potential interaction regions of the CheR_like_ domain, the residue sequence conservation using spirochete CheR_like_ domains was mapped onto the AlphaFold structure (using ConSurf with pre-aligned input sequences). The sequences used for this analysis are all the previously identified CheR_like_ domains, which are exclusive to spirochetes^29^. Overall, the CheR_like_ domain has high sequence conservation on two adjacent surface-exposed patches, which are both located on the C-terminal sub-domain (Fig. 5C). One conserved patch corresponds to the substrate-binding pocket, where the residues W256, G291, C292, E297, D321, H371, R388, and D389 are strictly conserved. The second patch is located on a ‘lid’ directly above the substrate pocket, where residues D323, L324, S328, F368, E369, Y370, H371, and D372 are strictly conserved (Fig. 5C). In the N-terminal sub-domain, there are five strictly conserved residues that are at least partially surface-exposed, but they do not form an obvious patch, and most of them are located at the interface with the C-terminal sub-domain. In total, ~14.5 % of residue in the CheR_like_ domain are strictly conserved (35/241 residues). Identical analyses were conducted using the AlphaFold *Td* classical CheR model and previously identified F2 spirochete CheR sequences^29^. In the classical CheR proteins, ~25.6 % of all residues are strictly conserved (69/269 residues). Conserved surface-exposed patches are located on areas that surround the buried substrate pocket, the N-terminal sub-domain, and the C-terminal region known to interact receptor substrates (Fig. S6B)^36^. Notably, the classical CheR does not possess the conserved, surface-exposed ‘lid’ above the substrate pocket since the ligand is buried (Fig. S6B). Additionally, the CheR_like_ domain is not conserved at the receptor binding regions of classical CheRs (Fig. 5C)^36^.

### The CheW protein domain is present in a diverse set of proteins

Given the role of CheWS in signal input, we investigated if the CheW protein domain is also present in other proteins besides the known CheV^43^, CheA^22^, 2xCheW^44^, 3xCheW^45^, and CheW-CZB^46^. Using the Pfam database we found 30 unique domain architectures that contain the CheW protein domain model that have not been described in the literature to date (Fig. 6). From this dataset, 12 were associated with some other domain proteins related to signal input according to the domain classification from the MiST3 database^47^.

**Fig. 6:**
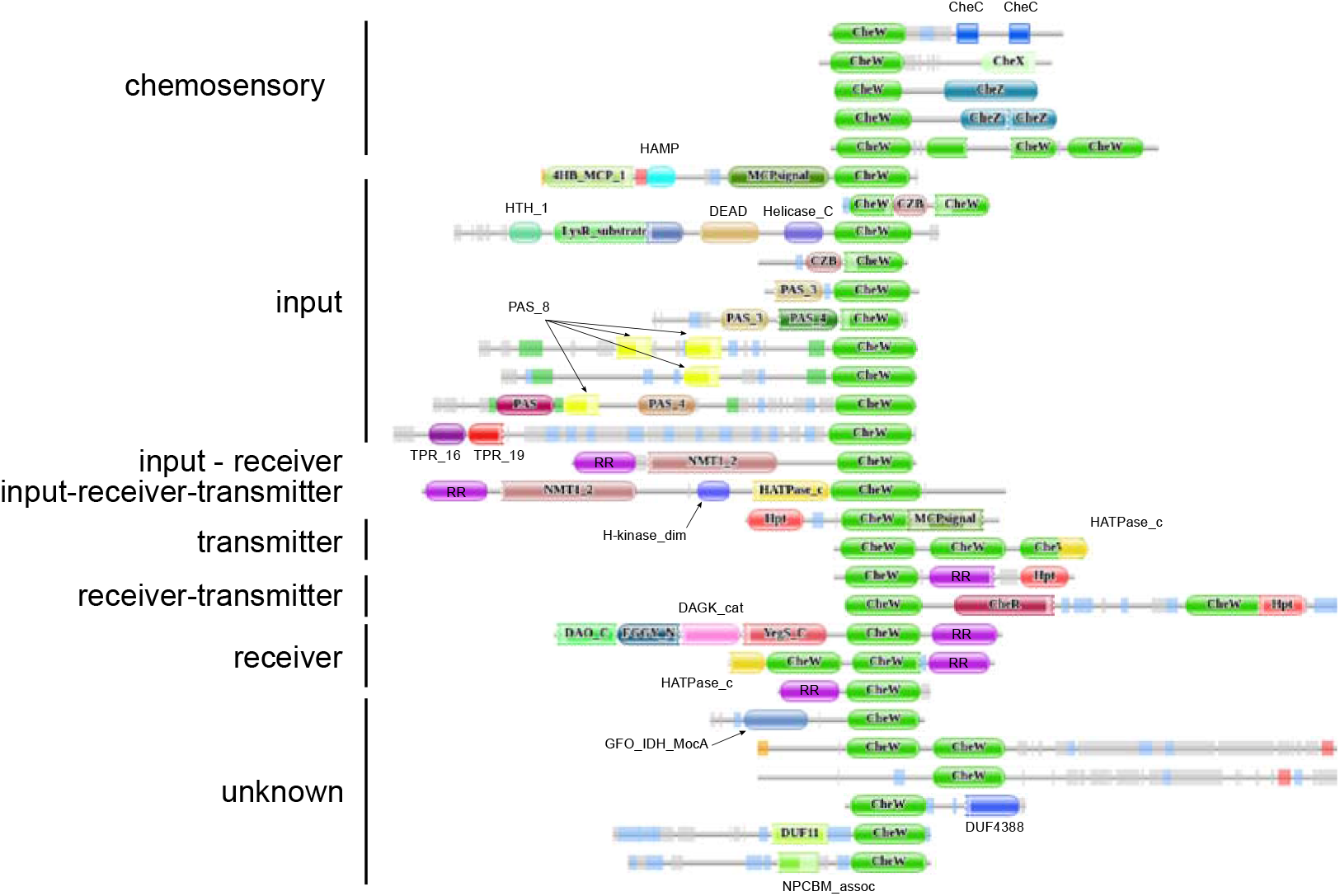
Protein domain architectures containing CheW domain model without a known function. The domain architectures are organized by their putative type of biological function. The RR domains are the Pfam Response_reg (PF00072). The architectures are annotated with predicted disordered (gray) and low complexity (blue) regions.

## Discussion

Here, we revised the model for round body formation in spirochetes, and established a link between round body formation and chemotaxis. An explanation for our observed deviation from the established model for round body formation could be that the previous visual characterizations utilized spirochete cells under environmental stress (reviewed in Bamm et al.)^10^, which increases round body prevalence. Additionally, these studies used traditional transmission or scanning electron microscopy techniques that require cell dehydration and/or chemical processing^48^. Cryo-ET, on the other hand, instantaneously freezes cells in vitreous ice without chemical treatment, reducing artifacts that result from cellular stress or chemical treatments^48^. Our results support the previous observation during stationary-phase growth, where nutrients are depleted and/or growth-inhibitory chemicals are accumulated. However, in log phase growth, where the cells are under more ideal metabolic conditions, we observed a not-yet described round body formation process. Under these conditions, round bodies are phenotypically distinctive and seemingly formed by a different mechanism. Log phase round bodies morphologically differ from established stress-induced round bodies in three aspects: they are significantly smaller, have a singular dense structure in the inner membrane, and do not contain spiral-shaped cells. These round bodies form by budding and pinching off from the cell, as opposed to forming a vacant outer membrane compartment seen during stationary phase^5,38^. Therefore, we propose that spirochetes in favorable conditions undergo a sporulation-like process to produce round bodies that resemble previously characterized ‘core structures’^12,13^. This process could be advantageous to spirochetes since core structures may stay dormant for several weeks^5,13^, thereby ensuring colony occupancy if conditions become suddenly unfavorable. If this hypothesis is true, pathogenic spirochetes are more virulent and persistent than originally thought – the cells are not just capable of protecting themselves from deleterious environments but they also continuously shed germination-delayed round bodies as a fail-safe for colony collapse. Notably, a previous study in *Bb* investigated the presence of round bodies in log phase and stationary phase cultures through traditional fluorescence microscopy^15^. This study reported that round bodies are not present during log phase. However, the images contain small spherical puncta that may be round bodies, but their small size make them difficult to convincingly detect them in these resolution limited light microscopy images.

Spirochete growth and peptidoglycan (PG) synthesis occurs at specific mid-cell sites where binary fission eventually ensues^49^. In contrast, the cell poles are regions where no cell wall expansion occurs^49^. Therefore, non-specific or uncontrolled blebbing at the cell tips due to PG growth is unlikely a determining factor for round body formation. In support of this, our cryo-ET experiments clearly demonstrate the presence of continuous PG layer around both the forming and disassociated log phase round bodies. As the log phase round bodies possess the PG layer and two membranes, they do not resemble outer-membrane vesicles (OMVs) that have been previously observed in spirochetes^50^.

We found that a specific protein that is associated with the bacterial chemotaxis system, called CheWS, serves as a functional link between chemotaxis and round body formation. Specifically, CheWS possesses a C-terminal CheR_like_ domain that functions as a SAM sensor to modulate these behaviors. In the *Td* WT strain, the presence of SAM decreases round body formation and increases biofilm formation. An exact opposite trend is produced when the CheR_like_ domain is absent in *Td*; round body formation increases and biofilm formation decreases. Collectively, these data indicate that the CheR_like_ domain is involved in physiological transitions that are modulated by SAM. Notably, samples that contain the WT strain without the addition of SAM do not produce identical results to the ΔCheR_like_ strain without SAM. These effects may be due to the fact the WT strain can still sense and respond to intracellular SAM levels that are sustained without exogenous SAM addition. However, interpretation of the ΔCheR_like_ experiments with SAM are less clear. Round body and biofilm quantification in the ΔCheR_like_ strain differs significantly when exogenous SAM is present. Therefore, the role of CheR_like_ in round body/biofilm formation could be more complex and involve additional interaction partners, or a separate system can still sense and respond to exogenous SAM. Nevertheless, it is clear that states linked to spirochete persistence are influenced by SAM and the CheR_like_ domain. Interestingly, a previous study shows that when *Td* is incubated with other oral pathogens, co-biofilms are more abundant and round body formation is reduced when compared to monospecies *Td* biofilms^51^. As the addition of exogenous SAM produces similar effects, it is possible that small molecules such as SAM may act as chemical messengers to facilitate these transitions and/or inter-microbial interactions. Indeed, oral pathogens are not found as monospecies *in vivo*, but as mixed biofilm populations that are metabolically symbiotic^52,53^.

Chemotaxis assays unambiguously demonstrate that the CheR_like_ domain is solely responsible for the SAM chemoattractant response; deletion of the domain eliminates a response to SAM specifically and entirely. It is known that motility and chemotaxis play a role in *Td* biofilm formation and cobiofilm formation^19,20^. Therefore, chemoattraction toward SAM may also be involved in the formation of SAM-induced biofilms. The CheR_like_ domain likely binds to SAM at a surface-exposed cleft on its C-terminal subdomain. The solvent exposure of the ligand-binding site may allow for free diffusion of SAM within the cell, thus allowing the domain to sense current intracellular SAM concentrations. To date, this is the first reported instance of a characterized CheW-fused chemotaxis sensor. Although the mechanism for signal dispersal by the CheR_like_ domain to the chemotaxis array is unclear, previous research has demonstrated that structural changes in CheW confer alterations to CheA activity^54,55^. These observations suggests that signals are propagated across chemotaxis arrays through CheW:CheA interactions. Therefore, CheR_like_ may impart SAM sensing through CheW-mediated modulation of CheA activity. Such interactions could occur through perturbations in the CheWS inter-domain linker or through direct contact of the two domains. Sequence conservation of the CheR_like_ domain reveals a strictly conserved patch adjacent to the SAM-binding pocket, indicating a plausible interface for protein interactions.

Interestingly, our bioinformatics analyses identified several classes of CheW proteins that possess additional fused domains. Two of these domains, CZB and PAS domains, are established sensor domains. CZB domains are known to act as hypochlorous acid (HOCl) sensors in a wide variety of proteins from diverse bacteria, and can regulate protein activities in an HOCl-dependent manner^46,56^. PAS domains present in eukaryotes and prokaryotes and bind various ligands and/or proteins to modulate functionally diverse sensory pathways^57^. Therefore, chemotactic sensing through CheW via a fused domain may be a wide-spread mechanism that allows bacteria to chemotactically respond to the current intracellular environment. Intriguingly, a recent report demonstrates that specific catecholamines are chemotactically sensed in *Vibrio campbellii* through direct binding to CheW^58^. Together, these data show that the previous designation of CheW as a linker protein is a partial misnomer. In addition to stabilizing chemotaxis arrays, CheW is responsible for transmitting signals across the arrays, and can function as an input site for ligand recognition. Due to these recent advancements, we propose to reclassify CheW as a ‘hub’ protein that interplays signals from multiple proteins and/or ligands to derive a complete and functional chemotaxis array.

The fact that spirochetes have evolved and maintained a system to sense SAM indicates that the ligand is critical for cell survival, either as a nutrient or an important environmental cue. The most well-studied pathogenic spirochete is *Bb*, which is a tick-borne obligate parasite. Several studies in *Bb* show that SAM-mediated metabolic reactions are essential for cell growth and survival^34^. Furthermore, small molecules that prevent these reactions have been identified as suitable antimicrobial agents for *Bb^59^.* In addition to being a necessary nutrient, metagenomics studies of tick microbiomes suggest that microbial SAM-mediated pathways may partly contribute to a more resilient microbiome^35^. Although analogous research in *Td* has not yet been conducted, SAM-mediated pathways are present in all spirochetes^34^. Determining if and how SAM is associated with *Td* pathogenesis in humans may greatly impact our current understanding of its virulence mechanisms.

In summary, we demonstrate an improved model for round body formation and identified a new class of protein sensor in spirochetes. Our results suggest that spirochetes may be more resilient than previously thought, as they continuously generate round bodies, even in favorable environments. A novel SAM-binding sensor, CheWS, influences round body formation, biofilm formation, and chemotaxis. Investigations into this system may reveal an alternative mechanism for chemotaxis signaling that occurs through CheW interactions and bypasses chemoreceptor-based inputs. Previous experiments have shown that some bacteria possess soluble chemotaxis arrays that are potentially responsible for sensing the current metabolic status of the cell^60^. However, cryo-ET experiments in *Td*^29^, Bb^61^, and *Tp*^62^ do not show the presence of such soluble arrays. Indeed, chemotactic sensing via CheW may be functionally similar to soluble arrays and allow cells to sense the intracellular environment. We also show that SAM decreases the presence of round bodies and increases biofilm formation in *Td*. Importantly, it has been demonstrated that oral spirochete infections acquire resilience through chemotaxis and synergistic communal biofilms^51,63^. Collectively, our results exemplify the importance of examining non-canonical sensory systems in pathogenic bacteria, identify SAM as an influential factor in the life-cycle of pathogenic spirochetes, and may offer new targets for antimicrobial therapies.

## Acknowledgements

We thank the Netherlands Centre for Electron Nanoscopy (NeCEN) for access to cryo-ET and cryo-EM data collection facilities. This work is part of the research program National Roadmap for Large-Scale Research Infrastructure 2017–2018 with project number 184.034.014, which is financed in part by the Dutch Research Council (NWO). This work was funded by a grant from the Dutch Research Council (NWO) Talent Program Veni grant awarded to A.R. Muok, and by grants from the National Institutes of Health awarded to C. Li: R01AI078958 and R01DE023080.

## Methods and Materials

### Bacterial strains and culture conditions

T. denticola ATCC 35405 (wild-type) was used in this study. Cells were grown in tryptone-yeast extract-gelatin-volatile fatty acids-serum (TYGVS) medium at 37°C in an anaerobic chamber in presence of 85% nitrogen, 10% carbon dioxide, and 5% hydrogen1. For biofilm assays, T. denticola cells were grown in oral bacterial growth medium (OBGM)2. T. denticola ΔCheR_like_ strain was grown with an appropriate antibiotic for selective pressure as needed: erythromycin (50 μg/ml)3. The ΔCheR_like_ strain was generated in a previous study3.

### Cryo-electron microscopy and tomography

T. denticola cells at log phase and stationary phase were concentrated 50X via centrifugation at 3000 xg for 5 minutes. For samples used for cryo-ET, protein A-treated 10 nm colloidal gold solution (Cell Microscopy Core, Utrecht University, The Netherlands) was added to a 1/10 dilution. For samples that contain S-adenosyl methionine (SAM), 0.5 mM SAM was added to the mixture and mixed by pipetting. For all samples, 3 μl aliquots of the mixture was applied to R2/2 200 mesh copper Quantiofoil grids (Quantifoil Micro Tools) that were freshly glow discharged. The grids were prepared using an automated Leica EM GP system (Leica Microsystems). The samples were pre-blotted for 60 s and blotted for 2 seconds in a chamber set at 95 % humidity and 20 °C. After blotting, the grids were immediately plunge frozen in liquid ethane.

Data collection for cryo-EM samples was conducted on a Talos transmission electron microscope operating at 120 kV. For cryo-ET experiments, data was collected on a Titan Krios transmission electron microscope (Thermo Fisher Scientific) operating at 300 kV equipped with a Gatan K3 Summit direct electron detector with a GIF Quantum energy filter (Gatan) at a slit width of 20 eV. Images were taken at a magnification of X26,000 which corresponds to a pixel size of 3.27 Å. Defocus was set to −6 μm and the total exposure was 100 e-/ Å2 per cell. Each tils series was collected with a bidirectional dose-symmetric tilt scheme with a 2° increment using Serial EM. Drift correcting and bread tracking-based tilt series alignment was done using IMOD4. Contrast transfer function determination and correction was done using CTF plotter5. Tomograms were reconstructive using simultaneous iterative reconstruction with binning set to 2 and iteration number set to 6.

### Quantification of round body formation

Log phase T. denticola cells undergoing round body formation were quantified via cryo-EM microscopy. Grids containing the WT strain or ΔCheRlike were prepared as described above and imaged on a Talos transmission electron microscope operating at 120 kV. For each sample, at least n = 113 cells were imaged and the percentage of cells undergoing round body formation was determined. For each strain, three separate cell cultures grown to the same OD_595_ were imaged and quantified with and without exogenous SAM added. The results are expressed as the mean percentage of cells forming round bodies +/- one standard deviation.

### Chemotaxis assays

The chemotaxis of T. denticola was tested by capillary assay3. Log-phase cultures of T. denticola were centrifuged at 5000 ×g for 7⍰min and supernatants were discarded. Cell pellets were resuspended in motility buffer (0.15⍰M NaCI, 10⍰mM NaH2PO4, 0.05⍰mM EDTA, 1% BSA, and 0.5% methylcellulose). The motility buffer was equilibrated in anaerobic chamber overnight. The final bacterial cell concentration was adjusted to 1⍰×⍰109cells/ml. Capillary tubes (0.025⍰mm inner diameter) were filled with either 0.5⍰mM hemin or 0.5⍰mM S-adenosylmethionine (SAM) (New England Biolabs, Ipswich, MA) in the motility buffer and sealed with vacuum silicone grease (Dow Corning, cat# Z273554-1EA) before they were inserted into bacterial suspensions (500⍰μl each). After incubation in anaerobic chamber at 37⍰°C for 2⍰h, the contents of each capillary tube were transferred to a new microcentrifuge tube and cell numbers were enumerated using a Petroff-Hausser counting chamber (Hausser Scientific, Horsham, PA). For the non-gradient control, capillary tubes were inserted into bacterial suspensions containing either 0.5⍰mM hemin or 0.5⍰mM SAM. The bacterial counts of each strain were normalized to those in the non-gradient control. Results are represented as the mean of cell numbers⍰±⍰one standard deviation.

### Biofilm formation assays

Biofilm formation was measured as previously described6 with slight modifications. Briefly, 200 μl of mid-logarithmic-phase T. denticola cultures (108 cells/ml) with or without 0.5 mM SAM was added into 96-well flat-bottom polystyrene plates. The plates were incubated anaerobically at 37°C for 7 days, allowing for biofilms to develop. The culture medium was carefully decanted, and the biofilms were stained with 150 μl of 0.1% crystal violet for 30 min, washed with water three times, and then air-dried. To quantify the amount of biofilms, 200 μl of 95% ethanol was added to each well and incubated for 30 min. The optical density at 595 nm (OD_595_) was measured using a Thermo Scientific™ Varioskan™ LUX multimode microplate reader (Bio-Rad). The results are represented as the average absorbance that is normalized to the WT strain without SAM addition ±⍰one standard deviation.

### Protein structural prediction and conservation analysis

The structure prediction of the T. denticola CheR_like_ domain from CheWS and the T. denticola classical CheR protein were done using the Colab notebook (dpmd.ai/alphafold-colab)7. Only residues 203-444 from CheWS were used, as these correspond to the CheR_like_ domain3. The resulting models were examined, aligned, and compared to existing structures using PyMOL. The PyMOL script ‘Color h’ was used to assess hydrophobicity of the models according to the Eisenberg hydrophobicity scale. Residue conservation analysis was conducted using ConSurf (https://consurf.tau.ac.il). The sequences used in the ConSurf analyses are all previously identified spirochete CheR_like_ domains and F2 classical CheR proteins3.

### Domain architecture diversity of sequences with at least one CheW

We used the web portal of Pfam database v35.08 to search for domain architectures containing the CheW Pfam model (PF01584). The database returns 111 unique domain architectures, including the single domain CheW protein. There were 75 domain architectures that also contain at least one HATPase_c (PF02518), Hpt (PF01627), or both domains, which together with the CheW domain, are strong markers for sequences of CheA9,10. We removed them from the analysis by conservatively considering them CheA homologs. There were 5 domain architectures attributed to proteins that have been identified before, including CheWS (CheW-CheRlike) in this study, that were also removed (CheV, 2XCheW, 3XCheW, CheW-CZB). To our knowledge, the remaining 30 architectures have yet to be studied and show the versatility of the CheW domains in proteins with a different biological functions. We classified these domain architectures according to their biological role using the categories in the MIST3 database list of signal transduction domains10. These architectures and the classification are shown in Figure 6.

---

**Fig S1:**
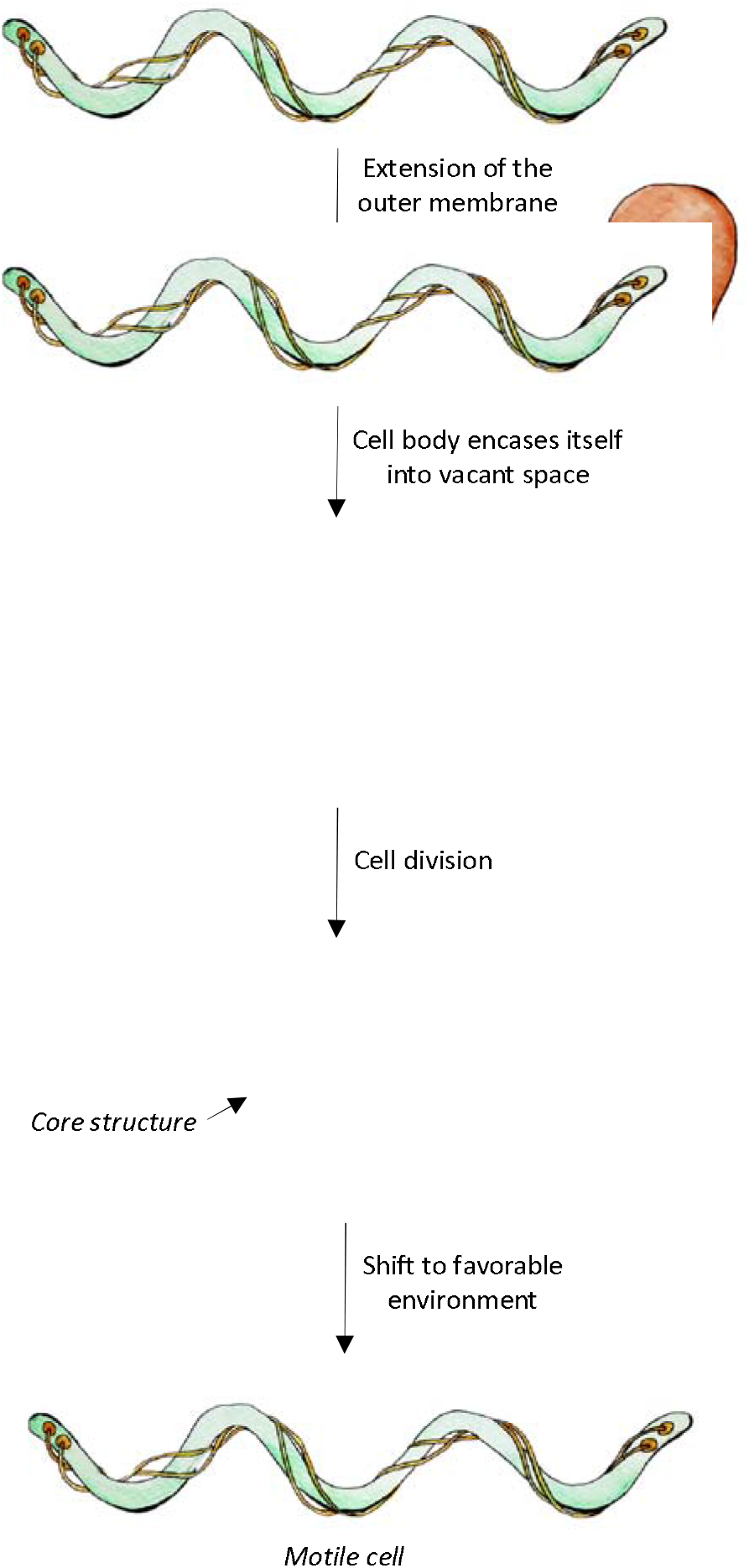
The established model for round body formation. Previous microscopy experiments in *Td, Tp*, and *Bb* have led to an established model for round body formation. In summary, unfavorable conditions induce the outer membrane to enlarge at the cell tip so that the spiral-shaped spirochete can move into the vacant space. Here, the cells are protected and can divide into more spiral cells or ‘core structures’ (dark blue). When the conditions are favorable, the cells are released and the spiral cells are motile. The core structures may take several weeks to form motile spiral cells.

**Fig. S2:**
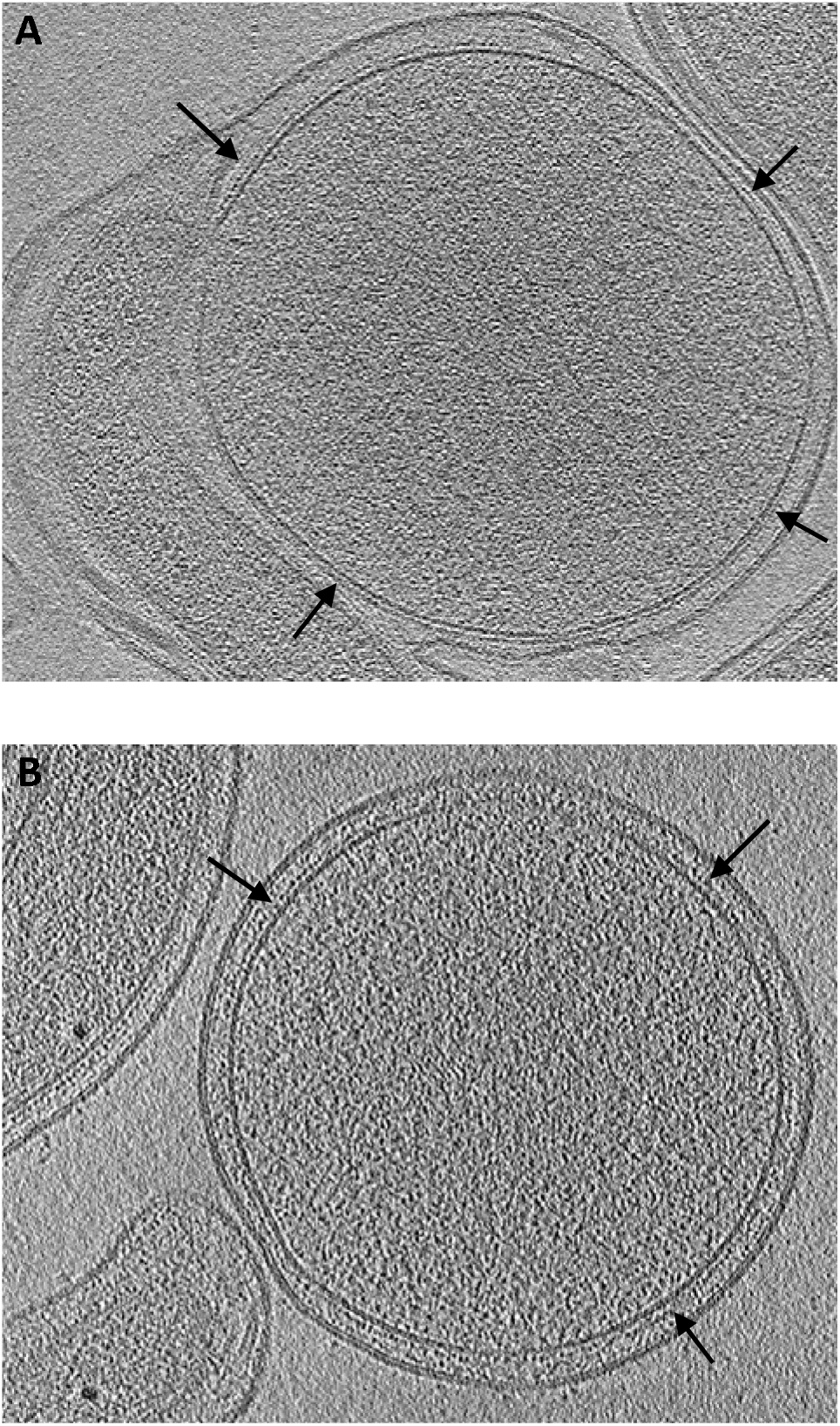
The peptidoglycan (PG) layer in log phase round bodies. **(A)** Log phase round bodies possess a continuous PG layer during their formation and **(B)** after separation from the spiral cells.

**Fig. S3:**
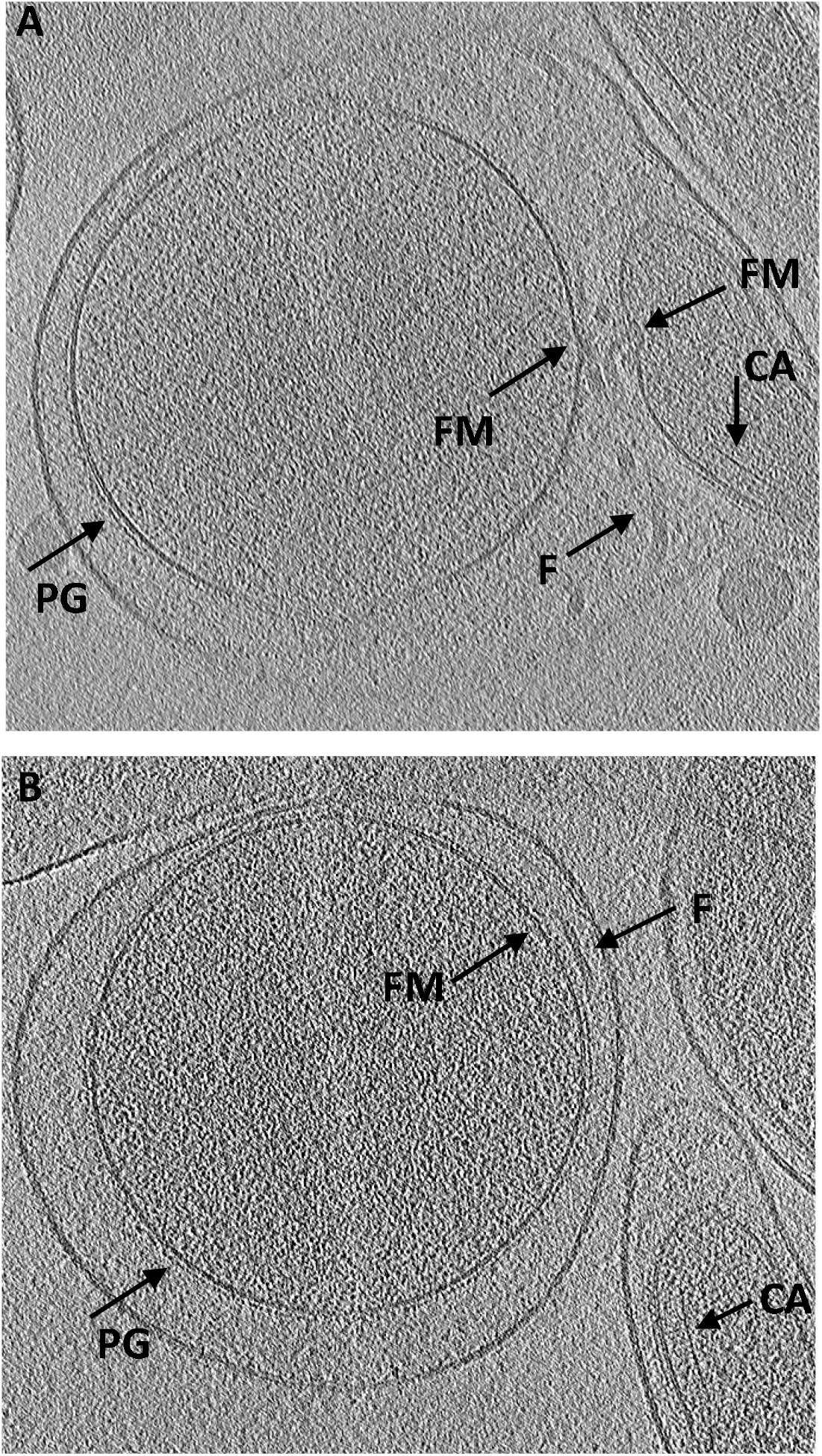
Motility machinery is present in log phase round bodies. **(A)** During their formation, flagella (F) and flagellar motors (FM) are attached to round bodies. The round bodies do not possess visible chemotaxis arrays (CA) but arrays are seen in the spiral cells near the forming round body. (B) After separation, round bodies can still possess flagella and flagellar motors.

**Fig. S4:**
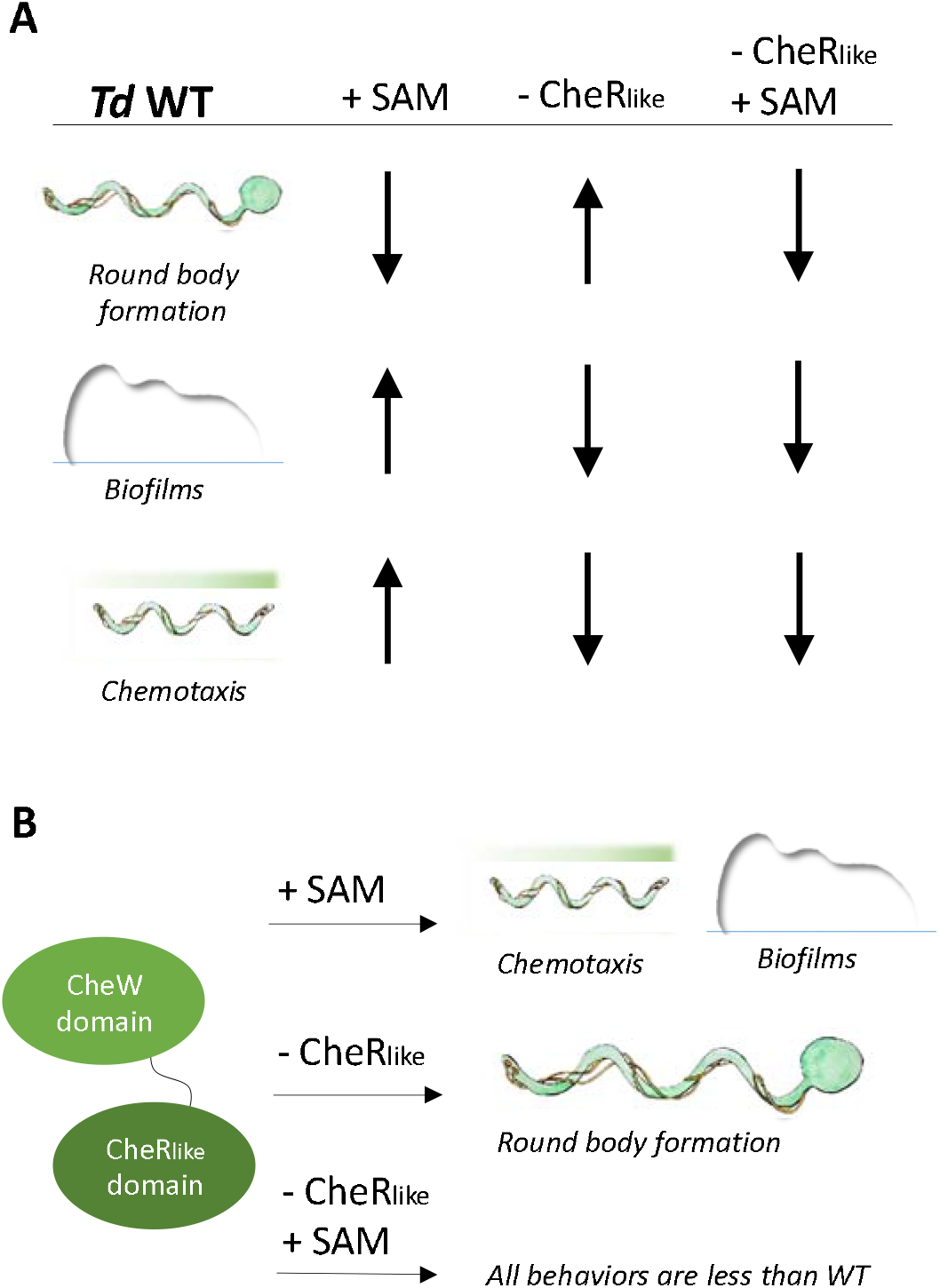
A graphical summary of the round body quantifications, biofilm quantifications, and chemotaxis assays. **(A)** Compared to WT cells without exogenous SAM added, the addition of SAM decreases round body formation, increases biofilm formation, and enables chemotaxis toward SAM. When the CheR_like_ domain is absent from the cells, the exact opposite trends are produced. When the CheR_like_ domain is absent and exogenous SAM is present, all behaviors are reduced when compared to WT without SAM. (B) The CheR_like_ domain enables chemotaxis toward SAM and biofilm formation. When the domain is absent, chemotaxis and biofilm formation are suppressed, but more cells undergo round body formation. When the domain is absent and SAM is present, all behaviors are relatively suppressed.

**Fig. S5:**
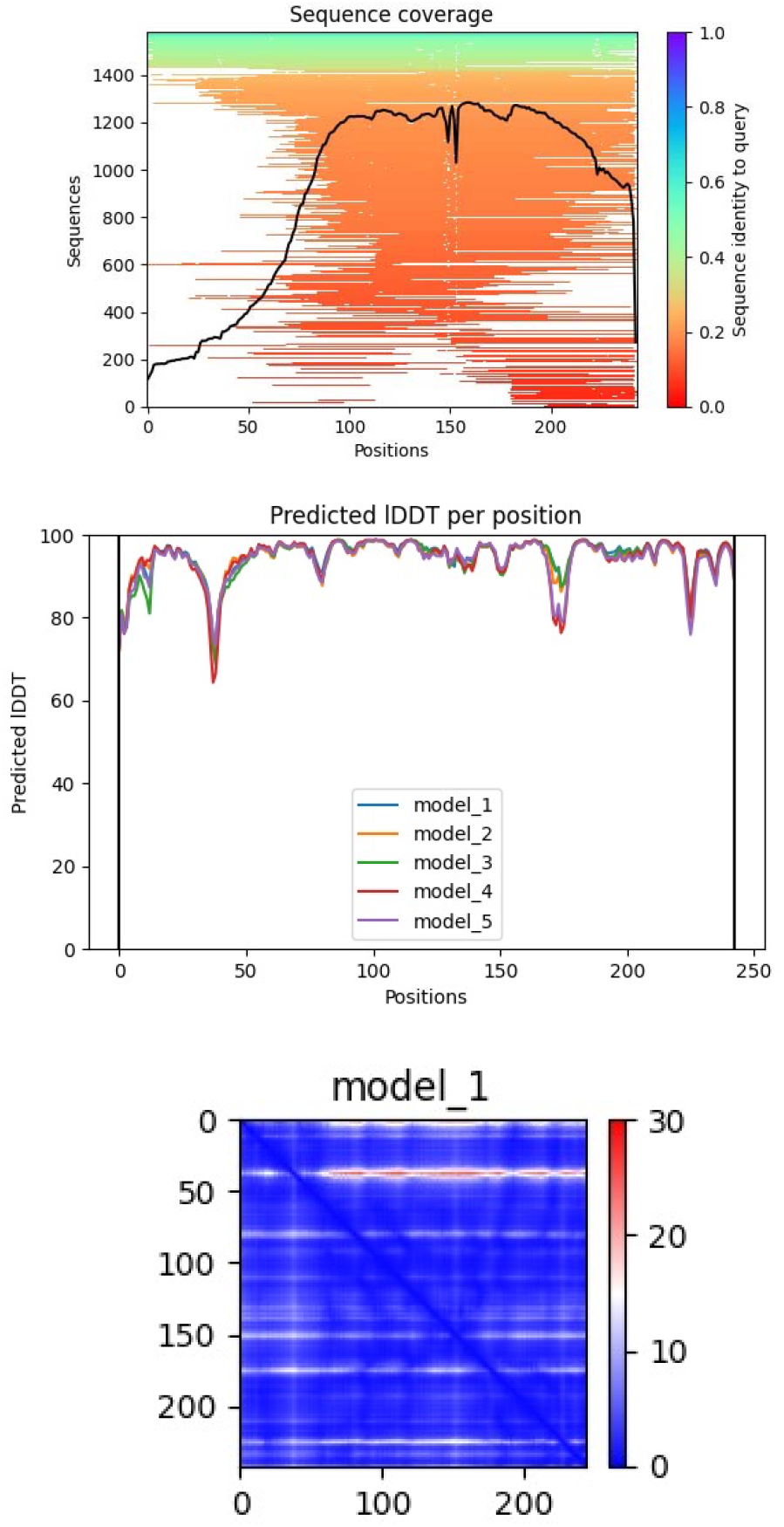
Output from AlphaFold 2.0 for the CheR_like_ domain model. Five models of the *Td* CheR_like_ domain were generated from the AlphaFold 2.0 Colab notebook (dpmd.ai/alphafold-colab). From the resulting models, Model 1 was chosen for further analysis.

**Fig. S6:**
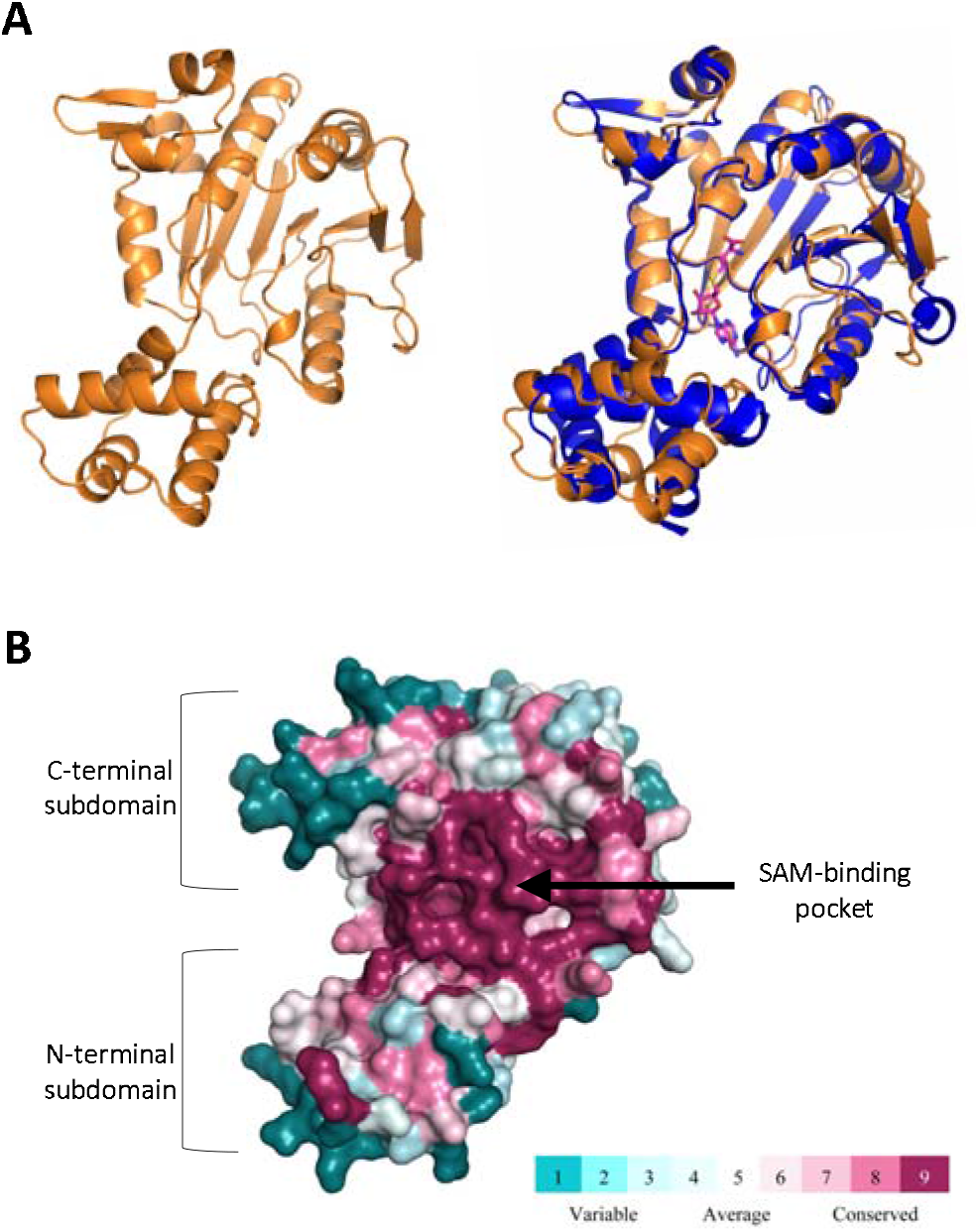
Structural analysis of the *Td* classical CheR protein model. **(A)** A model of the *Td* classical CheR protein was generated from the AlphaFold 2.0 Colab notebook (dpmd.ai/alphafold-colab). The model (orange) possesses the same topology and position of the subdomains as previously determined CheR structures (blue, PDBID: 5FTW), which buries the hypothetical SAM pocket. (B) Conservation analyses of the model reveals relatively high sequence conservation at the hypothetical SAM-biding pocket and regions known to interact with receptors.

